# A low-cost FPGA-based approach for pile-up corrected high-speed in vivo FLIM imaging

**DOI:** 10.1101/2024.12.17.628898

**Authors:** Felipe Velasquez Moros, Dorian Amiet, Rachel Megan Meister, Alexandra von Faber-Castell, Matthias Wyss, Aiman S. Saab, Paul Zbinden, Bruno Weber, Luca Ravotto

## Abstract

Intensity-based two-photon microscopy (2PM) is a cornerstone of biomedical research but lacks the ability to measure concentrations, a pivotal task for longitudinal studies and quantitative comparisons. Fluorescence Lifetime Imaging (FLIM) based on Time-Correlated Single Photon Counting (TCSPC) can overcome those limits but suffers from “pile-up” distortions at high photon count rates, severely limiting acquisition speed. We introduce the “laser period blind time” (LPBT) method to correct pile-up distortions in photon counting electronics, enabling reliable low-cost TCSPC-FLIM at high count rates. The correction was implemented on low-cost hardware based on a field programable gate array (FPGA) and validated using a combination of in silico simulations and in vitro, ex vivo and in vivo measurements. The LBPT approach achieves <3% error in lifetime measurements at count rates more than ten times higher than traditional limits, allowing robust FLIM imaging of sub-second metabolite dynamics with subcellular resolution. Our work enables high-precision, cost-effective FLIM imaging at rates comparable to commercial systems and at a fraction of the cost, facilitating the adoption of FLIM across all areas of research needing affordable, quantitative live imaging solutions.

## 1. Introduction

Two-photon excited fluorescence microscopy (2PM) has revolutionized biomedical research, allowing the investigation of biological dynamics with sub-cellular and sub-second resolution in living animals [1,2]. Fluorescent labeling with proteins and dyes has been used to elucidate the morphology and behavior of cells and blood vessels under physiological and pathological conditions, while functional readouts such as calcium imaging or blood flow measurements have shed light on dynamic phenomena such as neuronal activity and neurovascular coupling [3,4]. However, despite its successes, intensity-based 2PM is heavily affected by confounding factors such as fluorophore expression, tissue scattering and absorption, and sample motion. While these problems can be somewhat alleviated using ratiometric approaches, the wavelength dependence of scattering and absorption still prevents accurate quantitation. Thus, traditional 2PM is insufficient to investigate biological processes (such as metabolism) which require the precise quantification of molecule concentrations [5,6].

A more powerful approach is represented by the combination of 2PM with fluorescence lifetime imaging (FLIM), a microscopy technique based on measuring the average time delay between the excitation of a fluorophore and the subsequent emission of a photon. FLIM leverages on an intensive property of a fluorophore, the fluorescence lifetime, that does not depend on the number of detected photons, eliminating the intensity-related biases encountered in thick biological tissues [6]. Furthermore, while the signals obtained with intensiometric and ratiometric methods inherently depend on the detection apparatus, the fluorescence lifetime is an absolute property that can be associated to a concentration value and used to directly compare experiments performed on different samples or instruments and even by different laboratories. Thanks to these advantages, FLIM has been used in combination with fluorescent sensors to monitor pH, metabolites, small ions and the cellular environment in vivo [6,7], or as a label-free method to track cell metabolism [8,9] and perform tissue analysis in clinical settings [10,11].

Among the existing methods to measure fluorescence lifetime [6,12], time-correlated single photon counting (TCSPC) is particularly attractive for biomedical research as it can be seamlessly integrated in existing two-photon microscopes due to the common requirement of pulsed laser excitation. By recording the arrival time of every detected photon, TCSPC maximizes the information collected from an experiment while at the same time minimizing noise, providing the highest degree of accuracy and sensitivity among light detection methods [13,14]. Furthermore, modern TCSPC based on the time-tagged time-resolved (TTTR) scheme [15] offers the possibility to combine FLIM with other analysis methods such as fluorescence correlation spectroscopy (FCS), significantly increasing the capacity to resolve signals coming from different fluorophores and investigate dynamics over time scales spanning multiple orders of magnitude (ns to s) [16,17].

Nonetheless, TCSPC presents two main technical limitations, known as “dead time” and “pile-up”, which set an upper limit to the number of photons per unit time that can be accurately tagged [12,18]. Dead time refers to the time in which the time-tagging electronics are unable to detect a new photon after a previous detection, due to charging or discharging of capacitors or other resetting processes. Some detectors, such as single-photon avalanche diodes (SPAD), also exhibit a dead time while the avalanche is quenched until the bias voltage is restored [19].

Pile-up was first defined in literature as “…the possible detection - and loss - of a second photon in the same excitation pulse period as a previous one” [12]. This definition was coined to describe the distortion induced by timing systems that would only consider the first photon within an excitation cycle [20] (“classical” pile-up). While modern TCSPC systems do not exhibit this limitation, pile-up is still present due to the overlapping of the electrical pulses from the detector, resulting in a single threshold crossing for multiple photons. To avoid the intensity-dependent distortion of the fluorescence decay induced by dead time and pile-up, FLIM has traditionally been performed at detection count rates of ∼5% of the laser repetition rate [12]. While effective in solving the problem, this approach translates into significantly longer acquisition times, severely constraining the temporal scale of the biological processes that can be investigated.

Several methods have been proposed to compensate or correct the distortions generated by dead time and pile-up. In the case of classical pile-up, the number of counts in each bin can be corrected probabilistically [20]. Similarly, theoretical models have been presented to address electronics and detector dead times or pile-up in modern TCSPC. These models can then be used either to recursively compute the undistorted decay [21] or to modify the fitting model [22,23], accounting for the distortion within the loss function. Hardware-based methods have also been published, including matching the dead time of the detector to the laser repetition rate [24] and filtering out all cycles with more than one photon [25].

State-of-the-art approaches for pile-up correction have resulted in a leap in FLIM acquisition speed in recent years, opening new avenues in quantitative biological studies. Nevertheless, modifications of the fitting equations add complexity to the analysis, prevent the use of noise-robust non-fitting methods, and require instrument-specific calibrations. Hardware-based methods have the advantage of providing undistorted data directly, removing the need for time-consuming algorithms during the acquisition or processing of data. However, they either require (small area) detectors for which the dead time can be tuned [24] or expensive counting electronics that can accurately estimate the number of incident photons per cycle at any count rate [25]. Furthermore, state-of-the-art commercial FLIM electronics come at a considerable cost, especially when multichannel acquisition is required.

In this work, we demonstrate that pile-up distortions can be significantly alleviated with a simple correction algorithm implemented on low-cost (<5000 USD) FPGA-based electronics. This approach eliminates the need for instrumental calibrations, post-processing, or modifications to fitting algorithms. We provide a simple, affordable, and effective solution for high-speed FLIM on any multiphoton microscope, which will significantly facilitate its adoption not only in the biomedical field but across all areas of scientific research.

## 2. Materials and methods

### 2.1 Custom TCSPC electronics

The TCSPC acquisition device is based on the ZCU102 FPGA evaluation board. The main unit of this board is a Zynq UltraScale+™ XCZU9EG-2FFVB1156E MPSoC (multiprocessor system-on-chip) from AMD. Attached to the ZCU102 are two custom-made PCBs: one transfers the analog pulse signals from the single-photon detector and the laser trigger to differential digital signals, and the other contains level shifters to forward microscope synchronization signals such as frame changes, line starts and stops, and the pixel clock to the FPGA.

For TCSPC, it is crucial to accurately measure the time-of-arrival of both the laser trigger and detector pulses, which are, from an electrical point of view, short analog pulses. First, these pulse signals are transformed into two differential digital signals using high-speed comparators, plus an extra digital-to-analog converter (DAC) to set the threshold voltages. If the analog voltage of the inputs exceeds the threshold voltage, the comparator switches. Both digital comparator outputs are then forwarded to the FPGA where the (rising and/or falling) edges are individually time-stamped using two time-to-digital converters (TDCs). The TDCs themselves are implemented as taped-delay-line TDCs [26] in the FPGA following a previously reported implementation [27] while excluding the calibration steps (as the precision without calibration was deemed sufficient). Within the FPGA, the TDC outputs a stream of arrival times for both the laser and the detector. In this data stream, events that are detected within the time of one laser period from the detection of a previous event are deleted (LPBT correction). Then, by computing the time difference between detector arrival time to the most recent laser trigger, one gets the time between the photon detection and laser excitation (the so-called “relative time”). The microscope synchronization signals coming from the level-shifter board are timestamped in the same format as the photon-arrival events. Since these require less precision, they are just synchronized to the internal clock without using a TDC.

The combined and time-sorted information about photon and timing events is assembled in a binary data stream according to the “PT3” 32-bit TTTR format [15] and transmitted to a PC via TCP/IP using 1 Gbit/s Ethernet.

While the comparator and the TDC block on the FPGA can catch pulses with a minimum pulse width of 170 ps, the differential FPGA input might be more limited. Our tests showed that switching frequencies >1 GHz are possible, corresponding to a minimum pulse width <500 ps or a dead time (two consecutive pulses) of less than 1 ns. In terms of internal data processing speed, the TDC can handle >1000 Mcps for a very short time (around 300 pulses), while internal FPGA buffers overflow at constant count rates >200 Mcps. The theoretical maximum count rate forwarded to the event assignment block after discarding photons with our LPBT method is equal to the laser repetition rate of 80 Mcps. The assignment block can handle 200 million events per second in total, which is roughly 120 Mcps for the photon detection path at a laser repetition rate of 80 MHz. This means that when the LPBT method is enabled, buffer overflows in the assignment block are eliminated by design.

An additional buffer of 1 GB is present on the DDR memory to store assigned events in PT3 format (32 bits per event). At the maximum count rate of 80 Mcps, this buffer would fill at a speed of 320 MB/s. At the same time, transmission of the data through the 1 Gbit/s Ethernet connection would empty the buffer at a maximum speed of 125 MB/s (31 Mcps). Under these conditions (net rate of 195 MB/s), the buffer would overflow after about 5 seconds. However, as these very high count rates were achieved only in the brightest parts of our biological images, we never observed DDR buffer overflow in our experiments.

The accuracy of the electronics has been tested using a custom arbitrary waveform generator (AWG) containing a dual-channel 8 GSPS DAC and an FPGA that is used to generate a simulated data stream. On one output, the laser trigger is imitated as an 80 MHz square wave (50 samples maximum output, 50 samples minimum output). The photon stream on the other channels is imitated using triangular waves, whose linear rising edge can be timed with a precision of about 1 ps.

The timing jitter (38 ps RMS, 52 ps FWHM) was determined by analyzing the measured relative time from synthetic detector pulses with a constant relative time. This assumes the AWG’s contribution to the timing jitter is negligible or, conversely, results in a conservative estimate of the timing jitter of our electronics.

Then, the timing error *e* (5.8 ps) was determined as follows:

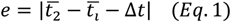

Where 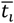 is the mean relative time of all detected events that correspond to *t* and similarly 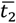 for events that correspond to *t* + Δ*t*.

### 2.2 TCSPC simulation

To test the performance of the LPBT method we implemented a simulation of the TCSPC data acquisition process. Every simulation included a total of 1000 molecules. For each molecule, using a time iteration process through every laser cycle within a fixed temporal window, a series of emission events was generated using random sampling based on a set emission probability threshold, and the time of the corresponding laser event recorded as “absolute time” (a typical convention in the TTTR format). The emission probability threshold was set in each experiment to match a specified average count rate (average photons per second). For each photon, the relative time was sampled from a probability distribution corresponding to the convolution of a monoexponential decay with a defined lifetime and a Gaussian instrument response function (IRF) with 200 ps FWHM. All single molecule traces were summed into a single trace representing the entire experiment. Sorting was performed to ensure the correct order of photon events and obtain the final TCSPC data in TTTR format. Finally, to ensure a consistent signal-to-noise ratio independently of the average count rate, for each simulation only the first 10^6^ photons were included in our analysis. The steps laid out above generate a realistic ground truth of synthetic TCSPC data. To simulate the TDC-based detection of voltage pulses, a continuous voltage trace was assembled by adding a predetermined single photon response (SPR) in correspondence with each photon arrival time. The photon events were tagged every time the voltage trace crossed a predefined threshold.

### 2.3 FLIM microscopy setup

A tunable-wavelength 80 MHz femtosecond laser (Chameleon Discovery NX with Total Power Control, Coherent) was sent into a custom two-photon microscope [28] and focused onto the sample through a water immersion objective. The fluorescence was collected through the same objective and split from the incoming laser light using a dichroic mirror (KS93 Cold Light Mirror, 685 nm LP, QiOptiq), then, the light was split by wavelength using a set of dichroic mirrors (F38-560, F38-506; AHF Analysentechnik) and filters (Brightline HC 542/50 or Brightline HC 475/50 depending on the channel, each combined with 785SP Edge, Semrock, USA) and detected using PMT detectors (H10770PA-40sel, Hamamatsu). To implement FLIM, one of the PMT detectors was replaced with a hybrid photomultiplier detector (PMA Hybrid 40 mod, PicoQuant) in combination with a tightly focusing lens (LB1761-A, Thorlabs) to minimize reflections.

The signal from the detector was fed into our custom TCSPC electronics alongside the 80 MHz synchronization signal provided by the laser. Furthermore, the frame and pixel clocks from the microscope scanning system (ScanImage Version 3.8 [29]) were also fed into the electronics such that they are integrated into the TCSPC data. This data was then transmitted to a PC via TCP/IP over 1Gbit/s Ethernet and processed using a C++ and MATLAB custom acquisition software, producing FLIM images containing a 256-bin decay for every pixel in the image.

### 2.4 Imaging of fluorescent solutions and IRF determination

To test our system under controlled conditions, we utilized sodium fluorescein solutions at different concentrations in pH 8 PBS. For unquenched fluorescein solutions, a concentration of 5×10^-6^ M was used. For solutions quenched with potassium iodide, the concentration was raised to 5×10^-5^ M to compensate for the loss of photons and allow to reach high count rates while minimizing excitation saturation effects. Imaging was performed at 950 nm excitation using a 20X water immersion objective (W Plan-Apochromat 20x/1.0, Zeiss).

To obtain measurements at different count rates, the power on the sample was varied using the integrated AOM in the laser. A beam sampler (BSF10-B, Thorlabs) was placed on the optical path to reflect a small percentage of light to a photodiode (PDA50B2, Thorlabs). By measuring the power under the objective with a power meter (S175C, Thorlabs), the voltage out of the photodiode can be used to determine the power on the sample during the FLIM measurements. For each excitation power, a measurement was made with and without the pileup correction enabled. To compute the lifetime of each measurement, a single decay (> 10^4^ photons) per image was calculated by subtracting the background image and summing over all the pixel-associated decays.

For in vivo imaging, the power at objective was maintained below 30 mW, while for slice experiments the power was kept below 150 mW.

IRF traces were measured by using second harmonic generation from KH_2_PO_4_ crystals. When necessary, background images were acquired using a water solution.

### 2.5 Fluorescence lifetime analysis

Exponential fitting was performed using a maximum-likelihood algorithm implemented in the FLIMfit 5.1.1 library [30], using either the FLIMfit GUI or using custom-written MATLAB wrapper code available at https://gitlab.com/einlabzurich/flimanalysis. For monoexponential fits, the lifetime value τ was extracted. For biexponential fits, the intensity-weighted lifetime τ*I* and the amplitude-weighted lifetime τ_*A*_ were computed as

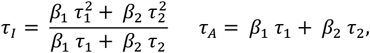

where τ_1_ and τ_2_ are the temporal constants of the two decay components and β_1_ and β_2_ are the respective fractional contributions to the initial intensity. The restricted average lifetime τ_8_ was computed using previously reported formulas [31].

The IRF temporal position was included as a fitting parameter to account for variations in optical path and periodic excitation was assumed to account for the incomplete decay of fluorescence at 80 MHz excitation frequency. When necessary, the background contribution was included in the fit. While pixel-wise exponential fitting is not feasible for the typical number of photons acquired in biological images, we performed global fitting, imposing identical values of τ_1_ and τ_2_ in all pixels of an image, unless averaging across regions-of-interest was performed before fitting.

The method-of-moments calculation was performed using the formula

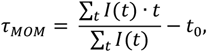

Where

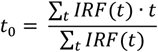

should be calculated as the center-of-mass of the IRF.

However, for lifetime components longer than approximately 1 ns this might give rise to imprecisions related to the incomplete decay at 80 MHz excitation. Since we were interested only in the variation of τ*MOM* within the same sample as function of count rate, we calculated *t*_0_ as the difference between τ and τ*MOM*at low count rates, where pile-up effects are negligible.

### 2.6 Animals

All experimental procedures were approved by the local veterinary authorities in Zurich and were conducted in accordance with the guidelines of the Swiss Animal Protection Law (Veterinary Office, Canton of Zurich; Act of Animal Protection, 16 December 2005, and Animal Protection Ordinance, 23 April 2008). For in vivo lactate measurements, one 5-month-old female wild-type mouse (C57BL/6J; Charles River) was used. For ex vivo imaging of ATP dynamics, one three-month-old female transgenic mouse C57BL/6J-Thy1.2-ATeam1.03^YEMK^ [32] was used. Mice were group-housed under standardized conditions (temperature: 20 ± 2°C, reversed 12-hour light/dark cycle) with ad libitum access to food and water.

### 2.7 Surgical interventions

Animals were anesthetized with a mixture of fentanyl (0.05 mg/kg body weight; Sintenyl, Sintetica), midazolam (5 mg/kg body weight; Dormicum, Roche), and medetomidine (0.5 mg/kg body weight; Domitor, Orion Pharma) injected subcutaneously. To prevent hypoxemia, 100% oxygen was administered via a face mask at a flow rate of 200 ml/min. Core temperature was maintained at 37°C using a homeothermic blanket heating system (Thermo Lux) throughout surgical and experimental procedures. Vitamin A ointment (VitA POS, Bausch+Lomb) was applied to the eyes to prevent corneal desiccation. Ringerfundin® (10 ml/kg body weight s.c.; B. Braun Medical AG) was administered for fluid replacement.

For head plate implantation, the animals' heads were secured with ear bars in a stereotaxic frame (Model 900; David Kopf Instruments), following established protocols [33]. Briefly, after removing head hair and disinfecting the scalp (Kodan® forte; Schülke & Mayr), local anesthetics (50 µl of stock solution consisting of lidocaine (1 ml, 20 mg/ml; Lidocain ad us. vet., Streuli) and bupivacaine (1 ml, 5 mg/ml; Bupivacain, Sintetica SA) mixed in saline (1 ml)) were injected subcutaneously into the incision area. A midline incision was then made from the eyes to the neck, and the underlying skull was carefully cleaned. A bonding agent (One Coat 7 universal; Coltene) was applied, followed by fixation of a custom-made metal head plate with light-curing dental composite (Tetric EvoFlow®, ivoclar vivadent).

A 3 × 3 mm craniotomy was performed using a dental drill (H-4-002 HP, Rotatec GmbH). Intracortical injections at 150 µm and 300 µm below the dura of adeno-associated virus (70 nl per injection spot, 4.7 x 10^11^ vg/mL of AAV6-hSYN1-chl-Lilac-QPRE-bGHp(A) (Viral Vector Facility, UZH) were conducted with a custom-made microinjector to achieve neuronal expression of the FLIM-based lactate sensor LiLac [34]. A square glass coverslip (3 x 3 mm; Powatec GmbH) was subsequently placed over the exposed brain surface and sealed with the light-curing dental composite.

Postoperative analgesia included buprenorphine (0.1 mg/kg body weight s.c.; Temgesic®, Indivior Schweiz AG) administered immediately after surgery, and carprofen (10 mg/kg body weight s.c.; Rimadyl Inj. ad us. vet., Pfizer) provided at 12-hour intervals.

### 2.8 In vivo imaging

Three weeks following virus injection, animals injected with the Lilac [34] sensor were imaged approximately 150-200 µm below the dura (cortical layers L2/3) at 2.96 Hz frame rate (pixel dwell time 3.2 ms) and 256×256 pixel resolution, using an excitation wavelength of 870 nm and a 25x water immersion objective (XLPlan N 25x/1.05w MP, WD = 2 mm, Olympus). During imaging, animals were head-fixed and anesthetized with the same anesthetic mixture as used for the surgical procedure (see “Surgical interventions”). Experimental protocol: Sodium L-lactate (L7022, Sigma-Aldrich) was administered intravenously via the tail vein over 3 min after a two-minute baseline acquisition. The injections were controlled by a peristaltic pump (Reglo digital ISM 831, Ismatec SA). Injections at dosages of 1.5 and 4 mmol/kg body weight were performed. FLIM images were motion-corrected and regions of interest corresponding to the somata of individual neurons were manually outlined using FIJI/ImageJ [35] prior to FLIM analysis. Analyzed data are presented as mean ± standard deviation (SD).

### 2.9 Brain slice preparation and imaging

Mice were deeply anesthetized with isoflurane and after decapitation, brains were quickly dissected in an ice-cold protective cutting solution (in mM: 135 NMDG, 1 KCl, 1.2 KH_2_PO_4_, 1.5 MgCl_2_, 0.5 CaCl_2_, 10 glucose, 20 choline bicarbonate, pH 7.4). Coronal sections (300 μm thick) were cut in the same solution using a vibratome (Vibration microtome, HM 650V, VWR) and afterwards kept in ACSF (in mM: 120 NaCl, 2.5 KCl, 1.25 NaH_2_PO_4_, 25 NaCO_3_, 1 MgCl_2_, 2 CaCl_2_, 10 glucose, 10 sucrose, 1 sodium lactate, 0.1 sodium pyruvate, pH 7.4) at 34°C for half an hour and then at room temperature until imaged. All solutions were continuously oxygenated with 95% O_2_ and 5% CO_2_.

Slices were transferred to a recording chamber (RC26, Warner Instruments) mounted on a temperature controlled Mini Bath Chamber (Luigs & Neumann) through an aluminum adapter and continuously perfused with oxygenated ACSF at a flow rate of 3 mL/min using a custom-based pumping system based on the PiFlow device [36]. Throughout all experiments, the temperature was maintained at 34°C. Slices were left in the recording chamber for at least 20 min before starting the experiment to ensure proper acclimatization.

Imaging of neuronal ATP dynamics was performed at 2.96 Hz frame rate (pixel dwell time 3.2 ms) and 256×256 pixel resolution, using an excitation wavelength of 870 nm and a 16× water immersion objective (N16×LWD-PF, 16×/0.8, Nikon). When necessary, spatial or temporal resolution reduction or region-of-interest averaging was performed in post-processing. For ATP depletion experiments, NaN_3_ (5 mM) was used to block mitochondrial respiration, as previously reported [37].

## 3. Results

### 3.1 The LPBT approach to pile-up correction

Threshold-based detection, in which the arrival time of a photon is determined by measuring when the voltage signal from a detector pulse crosses a certain predetermined value, is a simple and cost-effective photon counting method. Despite being widely used, threshold-based detection is affected by pile-up distortion. Every time a photon is detected, the voltage signal from the detector remains above the detection threshold for a certain “blind time” (dependent on the response time of the detector), preventing later photons from being detected (Figure 1a). Since the likelihood of two photons arriving close in time is proportional to the emission intensity, the loss is higher at shorter relative times, resulting in the typical distortion shown in Figure 1b. As the average incident photon rate increases, the effect of pile-up becomes progressively more significant.

**Figure 1.**
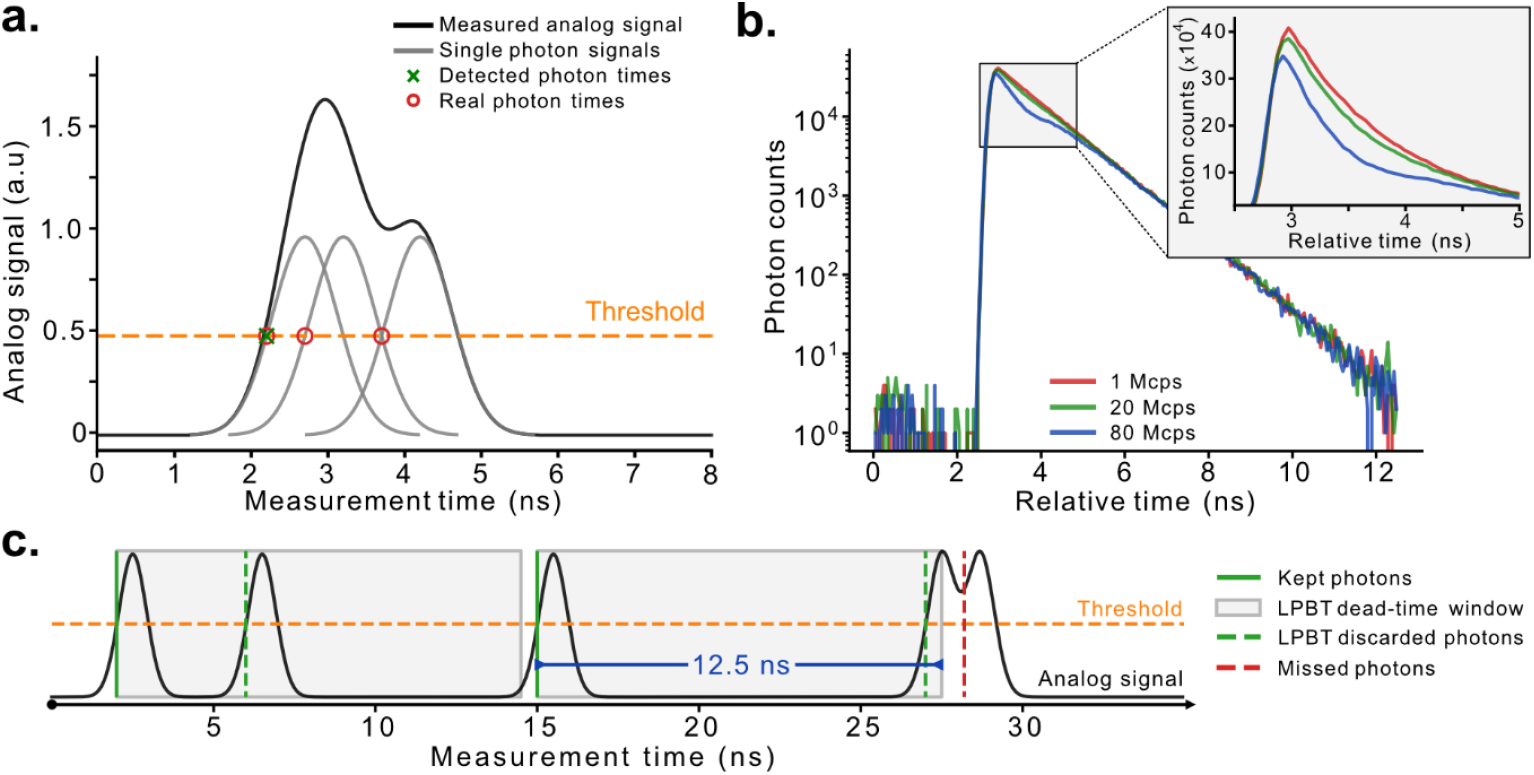
(a) Source of pile-up distortion in TCSPC measurements based on time-to-digital converters (TDCs): the analog signal from the detector is the sum of signals from individual photon events, resulting in a single threshold crossing when multiple photons reach the detector within a short time interval. (b) Impact of pile-up on a monoexponential fluorescence decay (1 ns lifetime, 10^6^ photons). The higher count rate decays have both a decreased brightness and distorted shape compared to the 1 megacount per second (Mcps) decay due to pile-up. (C) Working principle of the laser period blind time (LPBT) correction: after every detected photon (solid green lines) an artificial dead-time period of duration equal to the laser repetition rate is imposed (gray boxes) and photons that arrive during that period are discarded (dashed green lines). The real-world behavior of the LPBT correction is illustrated by the analog signal (black). Due to the finite pulse width of each photon event, a discarded photon can mask a valid photon that arrives soon after the artificial blind time period is over (dotted red line).

To counteract this effect, we implemented an approach, here defined as the “laser period blind time” (LPBT) method, that imposes an artificial blind time after every detected photon with a duration of exactly one excitation period. Intuitively, this method works by exploiting the periodic nature of the measurement to effectively re-enable acquisition at the same temporal shift with respect to the laser synchronization signal, albeit with one full period of delay (Figure 1c). This concept was initially proposed [24] and experimentally validated [19] for the correction of dead time effects in single-photon avalanche diode (SPAD) detectors but has never been applied to solve the pile-up problem in threshold-based TCSPC electronics.

We implement the LPBT approach on our FPGA-based electronics by simply disregarding any threshold crossing for a full excitation period after detecting a photon. However, shifting the blind time enforcement from the detector to the electronics introduces an additional source of error. As depicted in Figure 1c, when a photon is detected right before the end of the artificial blind time, the signal from the detector could still be above the threshold when the blind time ends, potentially preventing a “valid” photon from being detected.

To assess the impact of this non-ideality, we conducted numerical simulations that combined a Monte Carlo statistical generation of photon events with the application of a Gaussian single photon response (SPR) to each photon event, mimicking the signal generated from the detector. Monoexponential decays were generated for a series of lifetimes and count rates, while the total number of photons was kept constant in all cases. For each simulation, we evaluated the difference between the “ground truth” decay, obtained from all simulated events without any detection loss, and the decays obtained in the presence of pile-up and by applying the LPBT correction.

At first, to illustrate the accuracy of the correction for an ideal, infinitely sharp SPR, we applied it on the ground truth data by simply discarding the photons based on their known arrival times. As shown in Figure 2a, while the total number of detected photons decreases, the shape of the decay is maintained, indicating that photons were removed in an unbiased manner. Next, we evaluated the impact of the full width at half maximum (FWHM) of the SPR on the correction effectiveness, since it is at the origin of the non-ideality of our approach. As the pulse width increases, the accuracy of the correction decreases, providing unsatisfactory results for FWHM greater than 1 ns (Figure 2b). To evaluate the impact of the correction in a real-world scenario, we measured the SPR of the HPD detector used in this study (Supplementary Figure 1), which has a FWHM of about 0.7 ns. We set the threshold at about three quarters of the SPR peak height, a reasonable choice for experimental scenarios in which the aim is to find a good compromise between the reduction of the SPR width at threshold and the risk of losing photons due to the variability in pulse height from the detector.

**Figure 2.**
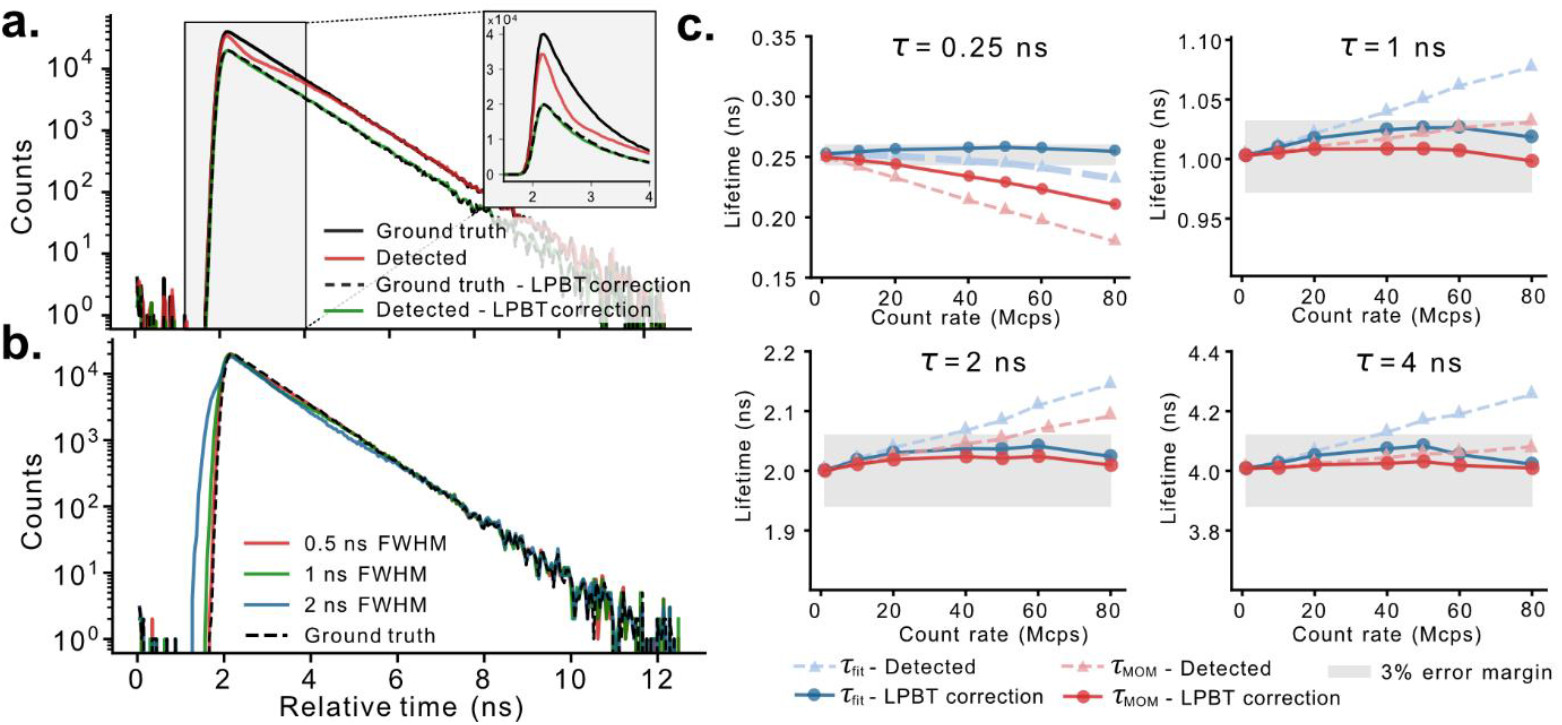
Validation of the LPBT correction with simulated data. (a) Fluorescence decays corresponding to the TCSPC data generated from one simulation (lifetime: 1 ns, count rate: 80 Mcps). The black lines correspond to the ground truth TCSPC data (i.e., no photons lost during measurement) including either all photons (solid line) or only the photons that are kept after applying the LPBT correction (dashed line). The colored lines correspond to the detected data resulting from tagging the threshold crossings of the simulated analog voltage trace including either all detected photons (red) or the photons that are kept after applying the LPBT correction (green). The inset shows a close-up of the region of the decay most affected by pile-up, in linear scale. (b) Effect of the FWHM of the SPR on the LPBT correction effectiveness (lifetime: 1 ns, count rate: 80 Mcps). (c) Maximum likelihood fitted and MOM lifetime estimates for the four decays from selected lifetimes (τ) and count rates. The gray shaded region illustrates the range of lifetimes that do not exceed 3% error compared to the ground truth. MOM lifetimes were shifted to match the fitted lifetimes at the lowest count rate (1 Mcps).

To evaluate the limits of our correction, we ran simulations using the HPD detector’s SPR and four different lifetime values, ranging from an ultrashort value of 250 ps to more realistic values of 1, 2 and 4 ns. For each lifetime value, simulations were conducted at progressively higher count rates. The upper value of 80 megacounts per second (Mcps), corresponding to an average of one photon per laser pulse, corresponds approximately to the maximum light intensity tolerated by our detector. Finally, the obtained decays (Supplementary Figure 2) were either fitted using a monoexponential model or estimated using the fit-free method-of-moments (MOM, also called “center-of-mass” method) which calculates the lifetime as a weighted average of the temporal decay [38,39]. As shown in Figure 2c, the uncorrected decay measurements significantly deviate from the ground truth across all tested lifetimes. However, after applying the LPBT correction, the calculated values fall within a 3% error margin from the ground truth for all simulated lifetimes and count rates, except for the MOM result for the shortest lifetime of 250 ps. While the deviation remains relatively small (0.02 ns at 40 Mcps), this suggests that with a combination of extremely short lifetimes and very high count rates, the accuracy of the correction may be reduced. Therefore, in these rare scenarios a careful characterization of the experimental system is recommended.

### 3.2 Hardware implementation and characterization

To streamline the adoption of FLIM across the bioimaging community, we designed a TCSPC system capable of handling the high count rates required to reliably study the dynamics of biological processes (tens of Mcps), without introducing pile-up distortions or requiring specialized algorithms to address them in post-processing. At the same time, to reduce cost and facilitate adoption, we aimed at compatibility with inexpensive electronics and common data transmission formats. To achieve these goals, we implemented a printed circuit board (PCB) based on a combination of comparators and digital-to-analog converters (DACs) to generate differential digital signals when the analog signals from the detector and the laser crossed a given threshold. A second PCB was designed to receive the scanning triggers from the microscope control electronics. The two boards were coupled to a time-to-digital converter (TDC) implemented on a commercial FPGA evaluation board to precisely measure the arrival time of photons and the laser synchronization signal, process the incoming data, and transmit it to an external PC via TCP/IP over Ethernet (Figure 3a).

**Figure 3.**
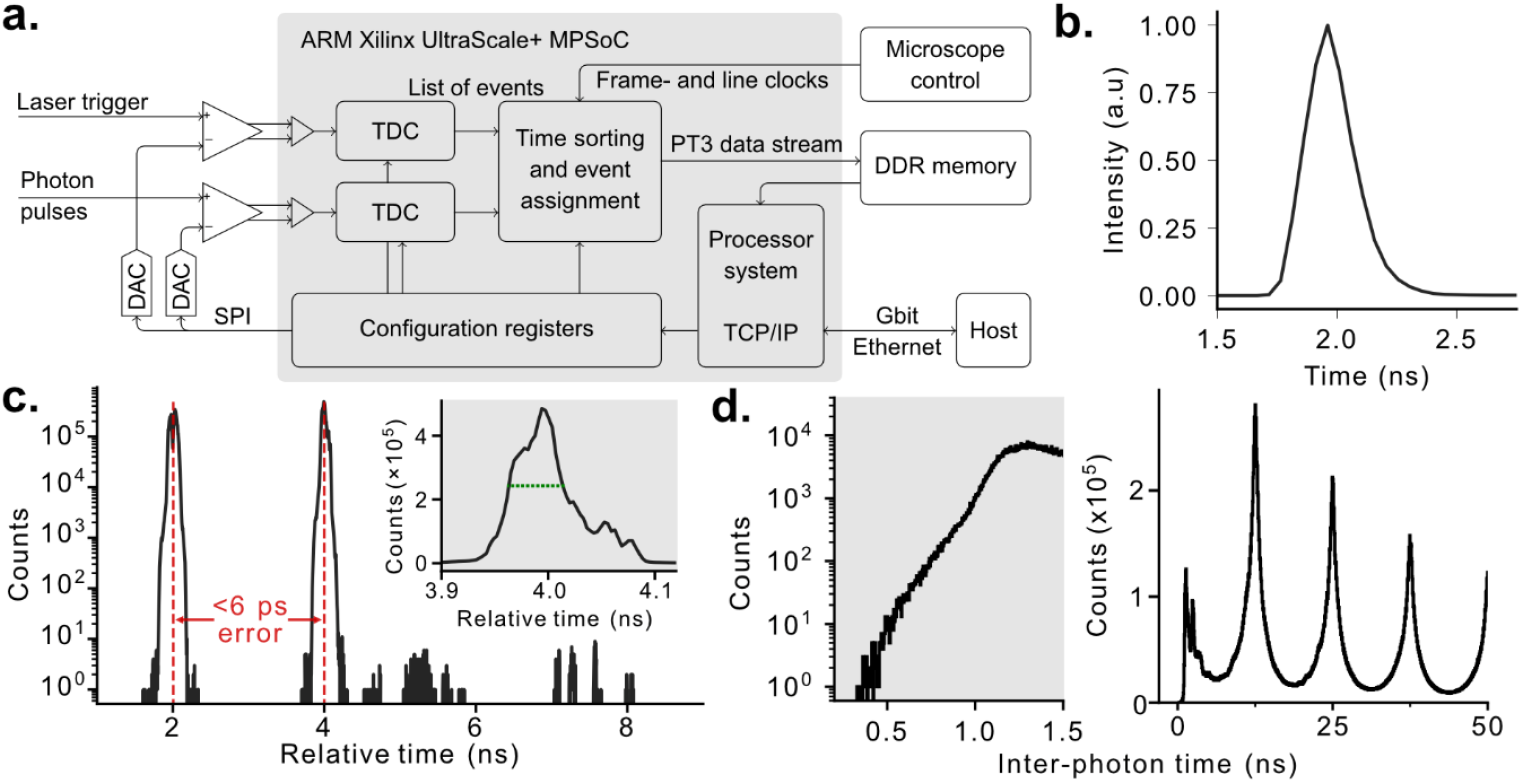
(a) Scheme of the custom electronics including the analog inputs from the laser trigger and photon detectors, the FPGA architecture, and additional connections required for the scanning synchronization clocks as well as for transmitting the data to a host PC. (b) Full system IRF acquired using the SHG signal from KH_2_PO_4_ crystals. (c) Timing performance of the custom TCSPC electronics measured using narrow voltage pulses separated by 2 ns. The inset shows a close-up of the temporal response in correspondence with the pulse at 4 ns. Inter-photon time histogram obtained from an IRF measurement at a high count rate. Left: close-up of the time interval between 0 and 1.5 ns; Right: a longer time range showing multiple laser cycles.

To assess the timing performance of our device, we used a custom-built arbitrary waveform generator (AWG) configured to generate one voltage pulse every excitation cycle. The relative time for each pulse was randomly assigned to be 2 ns or 4 ns, and the maximal temporal resolution of the TDC (3.26 ps) was used (Figure 3c). Our electronics was able to retrieve the correct distribution of incoming pulses with a timing jitter of 52.1 ps (FWHM) and a timing error for the separation between the two pulses of 5.8 ps. Using a laser repetition rate of 80 MHz and 256 temporal bins (an appropriate number for FLIM measurements [40]), the temporal duration of each bin is 48.8 ps. Under these conditions, the impact of the jitter of the counting electronics would barely affect the decay. Nevertheless, the temporal jitter of the whole FLIM system, the so-called instrument response function (IRF), was found to be about 220 ps (Figure 3b). Since the IRF compounds the inaccuracies of the timing electronics with the timing jitter of the detector and laser synchronization signals, the latter are clearly the dominant factors, and thus the precision of our electronics is completely adequate.

The blind time of our system can be determined by analyzing the histogram of the time intervals between consecutive photons, or inter-photon times (Figure 3d). The histogram exhibits a sharp decline at around 1.1 ns, while the absolute temporal limit appears to be about 0.5 ns. The former value is determined by the pile-up effect (two finite SPR pulses merging into one), while the latter is the intrinsic deadtime of the TDC. While theoretically it would not be possible to observe photons arriving at inter-photon times shorter than the limit determined by pile-up, their presence can be explained by variability in the pulse heights received from the detectors. Pulses with a smaller height have a very short pulse width at threshold voltage, allowing detection of a following pulse much earlier than for the average pulse height.

### 3.3 In vitro and ex vivo validation of the LPBT correction

The capacity of our approach to retrieve correct lifetime decays at different count rates was first evaluated in vitro using fluorescein solutions (5×10^-6^ M in PBS, pH 8). When the measurements were conducted at low count rates (<5 Mcps), a monoexponential fit returned the expected lifetime value of 4.1 ns [41], validating the accuracy of our electronics. However, when the average count rate was progressively increased by increasing the excitation power, a significant distortion of the decay was observed, with a corresponding deviation of the calculated lifetime (Figure 4a). Since at any given count rate the pile-up distortion becomes increasingly severe the shorter the lifetime, we utilized potassium iodide as a quencher to produce solutions with lifetimes of approximately 0.5 ns and 2 ns.

**Figure 4.**
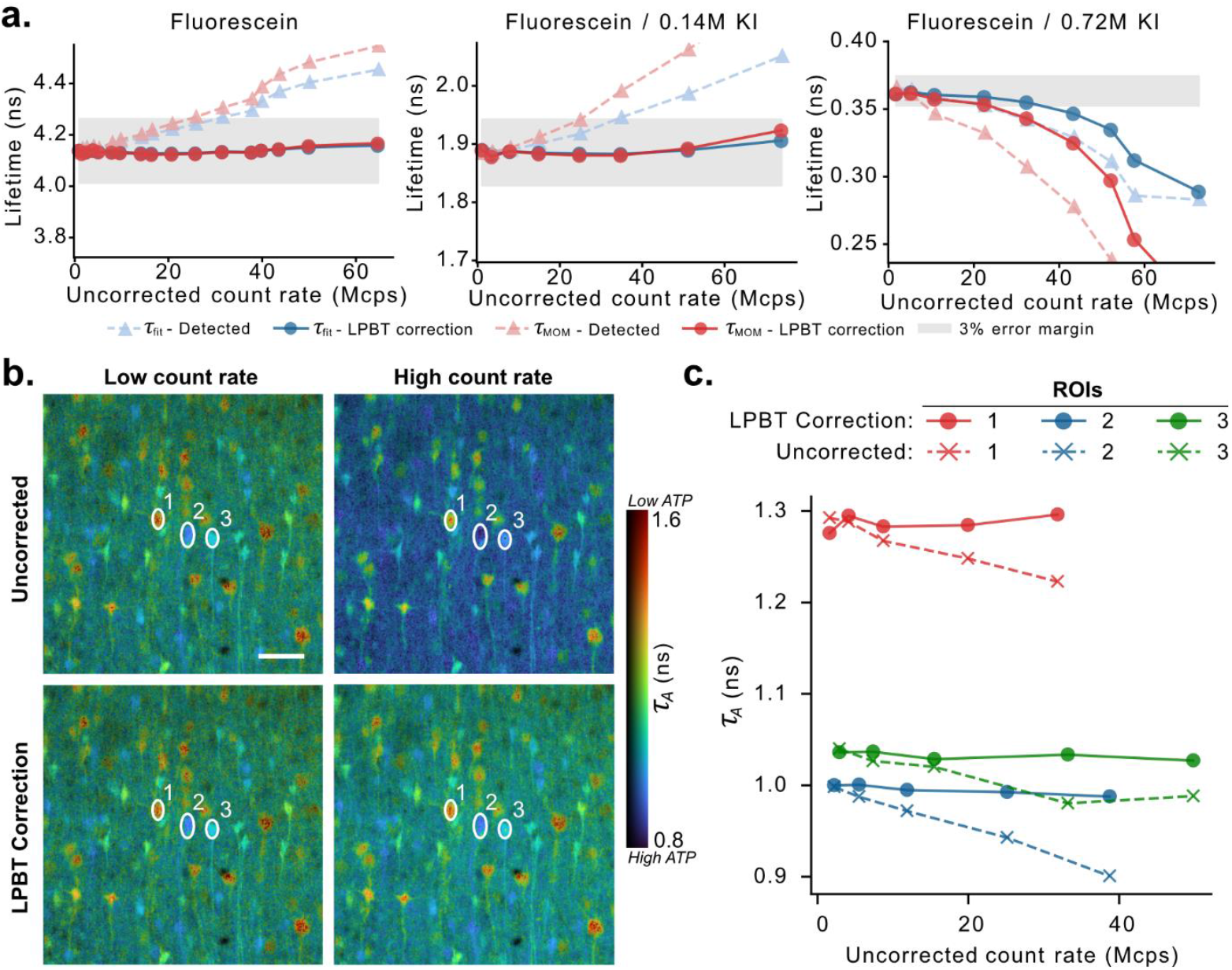
(a) τ (blue) and τ_MOM_(red) lifetimes calculated for the different KI-quenched fluorescein solutions tested at a series of excitation powers, before (triangles) or after (circles) application of the LPBT correction. The detected count rate without the LPBT correction is used for the x axis. The shaded regions highlight a 3% error margin around the fitted lifetime values measured at low count rates (<5 Mcps). The three panels reflect measurements done in absence (left) or progressively higher concentrations of KI (center and right). The τ_MOM_values are shifted to match the τ values at low count rates. (b) FLIM images (τ_A_) of the neuron-targeted ATP sensor ATeam1.03YEMK in mouse cortical slices at different count rates before (top row) and after (bottom row) applying the LPBT correction. The ‘low count rate’ images (first column) have average and maximum detected count rates of 1 and 6 Mcps, respectively. The ‘high count rate’ images have average and maximum detected count rates of 19 and 85 Mcps, respectively. (c) The change in τ_A_for each of the three selected ROIs displayed in (b) as the detected count rate increases. Scale bar = 40 μm.

At all lifetimes, the LPBT correction resulted in lifetimes estimates with errors below 3% for count rates of more than 65 Mcps (the maximum count rate at all lifetimes was dictated by the triggering of the overload protection of our HPD detector in a linearly spaced excitation power sweep), except for the short lifetime curves, which present significant deviations above 30-40 Mcps. In these conditions, additional distortions of the decay were observed, likely originating from operating the detector beyond the recommended maximum count rates for such a short lifetime. Similar trends were observed analyzing the decay using the MOM.

To demonstrate the applicability of our approach in biological imaging, we performed measurements on ex vivo mouse brain slices expressing the ATP sensor ATeam1.03YEMK in neurons. Under baseline conditions, we acquired images at progressively higher excitation powers, thus increasing the detection rate. By controlling the acquisition time, we ensured that all images had a similar number of total detected photons, so that any difference in lifetime cannot be ascribed to noise-dependent numerical effects.

Biological FLIM experiments are typically analyzed using a biexponential fitting model, out of which an average lifetime (either the “amplitude weighted lifetime” τ_*A*_ or the “intensity weighted lifetime” τ_*I*_ [38]) is extracted. However, under the typical number of photons per pixel acquired in biological imaging (hundreds to thousands), a pixel-by-pixel analysis would be unfeasible due to an insufficient signal-to-noise ratio (SNR). To overcome this obstacle, the analysis can be conducted on the average decay from a specific region-of-interest (ROI), or using global fitting under the assumption that the two decay constants of the model have identical values in every pixel [30,42]. Alternatively, the MOM provides a more noise-robust way to estimate an average lifetime (theoretically equivalent to τ_*I*_ [38]) on a pixel-by-pixel basis.

As visible in Figure 4b, uncorrected images show intensity-dependent changes in τ_*A*_, while after application of our LPBT approach the lifetime values were independent from the detected count rate. A ROI-based analysis on selected cells (Figure 4c) highlights the count-rate dependent bias and its correction by our approach. Similar conclusions can be drawn for τ_*I*_ and MOM results, although the drifts introduced a positive rather than a negative bias (Supplementary Figure 3).

### 3.4 Ex vivo and in vivo FLIM imaging of fast metabolite dynamics

To demonstrate the capacity of the system to acquire valuable FLIM information at high count rates, we performed an experiment in which ATP levels were depleted by application of the mitochondrial respiration blocker NaN_3_ (Figure 5a and b). Images were acquired continuously for 50 seconds every minute (the 10 seconds intervals were added due to memory management limitations of the current version of our acquisition software). During the continuous imaging periods, images were acquired using a frame rate of approximately 3 Hz and a resolution of 256×256 pixels. After LPBT correction, the maximum count rate reached in the experiment was 44 Mcps, while average count rates oscillated between 14 and 17 Mcps, resulting in an average of about 50 and a maximum of about 150 photons per pixel.

**Figure 5.**
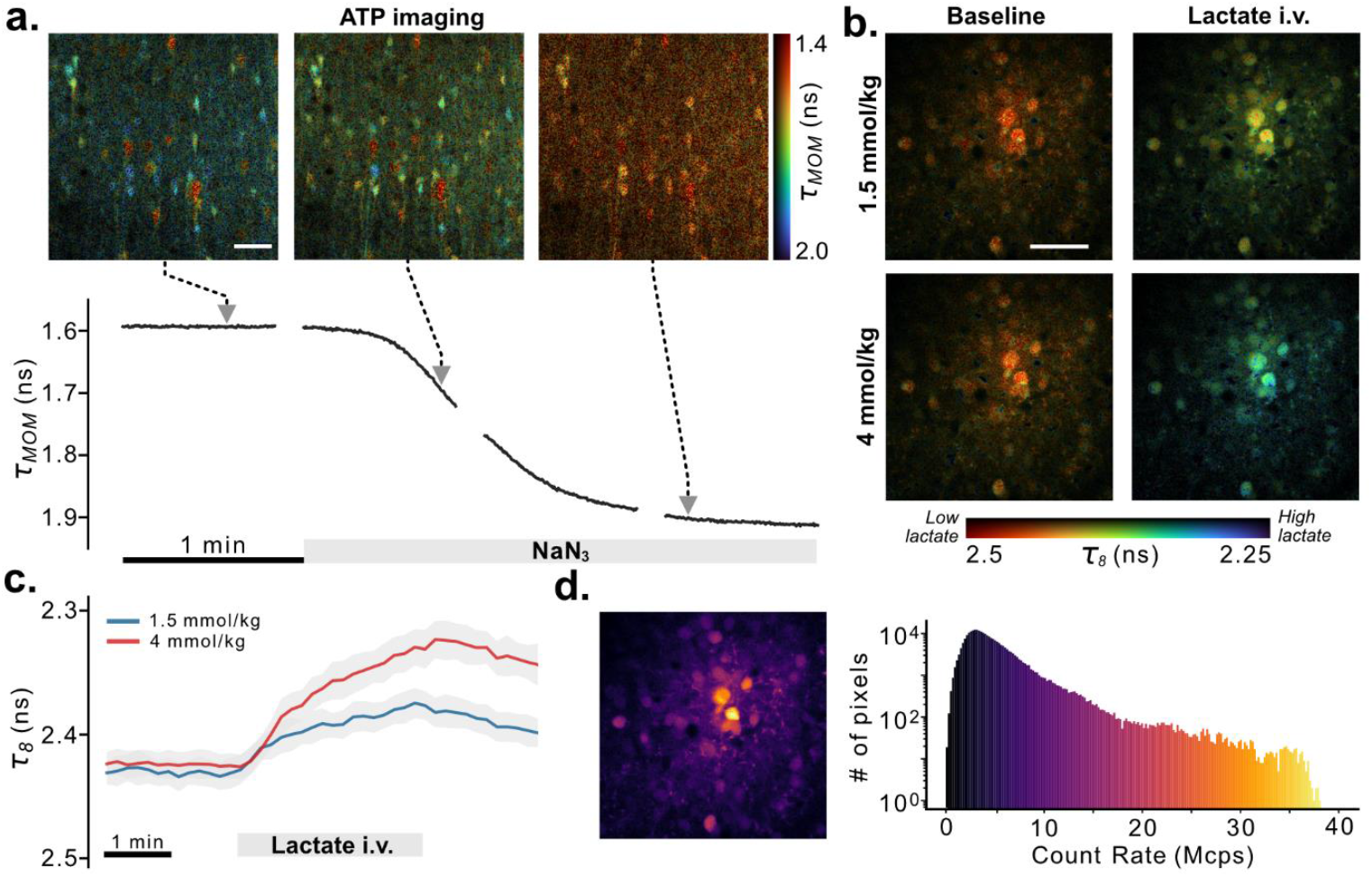
(a) Top: FLIM images showing different time points of the response of the neuron-targeted ATP sensor ATeam1.03YEMK to the application of NaN_3_ in mouse cortical slices. Each image shows the MOM lifetime estimate from the sum of 3 adjacent frames (1 Hz effective frame rate). Bottom: Frame-averaged response curve at 3 Hz. (b) FLIM images showing the response of the neuron-targeted lactate sensor Lilac in the somatosensory cortex of anesthetized mice to different concentrations of i.v.-injected L-lactate. (c) Lifetime dynamics of Lilac upon i.v. injection of L-lactate. Each curve shows the mean and SD of ROI-based curves corresponding to single neurons. (d) Color-coded image and histogram showing the large distribution of count rates in the images in (b). Scale bar = 40 μm.

Under these SNR conditions, we first used the initial baseline recording to determine if our analysis methods could provide reliable spatiotemporal lifetime information. For all methods, we compared the frame-averaged value of each lifetime estimator obtained on single frames with a ground-truth obtained from summing all the 150 frames of the baseline before analysis (Supplementary Figure 4). To assess the effect of reducing the spatial or temporal resolution, additional analysis was performed in temporally (1 Hz) or spatially (128×128 pixels) binned images. For τ_*A*_, spatial or temporal binning resulted in a detectable, although very small, change in the frame-averaged value, while for τ_*I*_ and MOM, no difference was detected in any condition. On the other hand, visual inspection of the images (Supplementary Figures 5-6) shows that, for all methods, the finest spatial features (such as neuronal processes) might be difficult to identify at 3 Hz and 256×256 resolution, requiring a decrease of the temporal resolution. Alternatively, the spatial resolution could be reduced, provided that this is accompanied by an appropriate reduction of the field-of-view to account for the loss of spatial detail.

As shown in Figure 5 and Supplementary Videos 1-2, the combination of our correction method with the MOM provides a convenient and robust method to image the evolution of concentrations in biological systems on a second and sub-second scale.

Finally, to further demonstrate the utility of our method for the characterization of in vivo dynamics of metabolites, we imaged lactate dynamics in the somatosensory cortex of anesthetized mice expressing the lactate sensor Lilac [34] in neurons (Figures 5b,c) with a temporal resolution of 10 seconds between images. The temporal resolution was chosen so that every single image, containing thousands of photons per pixel in the neuronal areas, could be analyzed with a high degree of accuracy and spatial detail. These imaging conditions are typical of slow metabolic processes for which it is often convenient to sum all collected frames in a single FLIM image, thus sacrificing temporal resolution to avoid generating an unnecessarily large amount of data. Upon an i.v. bolus injection of L-lactate, a clear rise is observed, followed by the onset of a decrease after the end of the injection (Figure 5c). As expected from an undistorted FLIM experiment, the lifetime response was dependent on the amount of injected lactate but not on the brightness (Figure 5d) of each individual cell. These results demonstrate the robustness of our approach in imaging fast changes of metabolites, even in (common) cases in which the chosen field-of-view presents large variability in terms of brightness.

## 4. Discussion

Fluorescence lifetime imaging (FLIM) has the potential to foster biomedical research by enabling robust, quantitative concentration measurements across samples. The quantitative power offered by FLIM enables the direct comparison of biomolecule levels between different experimental conditions, disease models, or even species while reducing the number of control experiments to rule out artifacts (such as the use of non-responsive sensors or separate tests to establish a sensor’s minimum or saturation levels). Despite these advantages, the widespread adoption of FLIM has been constrained by the lack of straightforward approaches to correct pile-up distortions that would otherwise limit its application for capturing biological dynamics on second and sub-second timescales.

In this work, we describe a simple and affordable strategy to eliminate distortions originating from pulse pile-up in TCSPC. While the mathematical principle of the correction was originally presented to compensate for dead time effects in custom-made SPAD detectors [19,24], we demonstrated for the first time its application to the correction of pile-up effects in threshold-based detection electronics. Although our implementation is an empirical approximation of the mathematically exact case, the correction virtually eliminates decay distortions when used in conjunction with detectors featuring a sub-ns FWHM single photon response. While this requirement excludes the use of PMT or SiPM detectors, it applies to HPD detectors, which are the most common choice for FLIM in living animals anyways due to their large active area, low noise, and high dynamic range.

Our method works with a maximum theoretical collection rate equal to the laser repetition rate, thus limiting the maximum SNR achievable for a given acquisition time, a problem that is exacerbated for lower repetition rate lasers. Nevertheless, for the typical 80 MHz laser used in two-photon imaging, the main limitation we encountered was the protection circuit of our detector (about 50 Mcps after LPBT correction). Under those conditions, we have demonstrated that undistorted FLIM imaging can be achieved with subcellular and sub-second resolution.

Using our approach, we implemented a TDC-based photon-counting device for a total price of <5000 USD in which all pile-up distortions are addressed in hardware using a lifetime-independent method. In the current version, the device is capable only of single-channel acquisition. While this is sufficient for many applications, it would be possible to implement several simultaneous TCSPC channels at a similar cost, as the FPGA is not operating at maximum capacity (or alternatively, use a cheaper FPGA while maintaining the same performance). Compared to the state-of-the-art, our strategy presents the major advantages of not requiring expensive electronics based on analog-to-digital converters (ADCs) or additional modifications to the fitting equations. The latter factor is particularly important, as it eliminates the need for instrument-specific calibrations and extends the scope of the correction to fast lifetime estimation methods, such as the MOM or phasor plots [6].

This unique advantage offers the possibility to implement the MOM method directly on the FPGA, computing and transmitting the FLIM results in real time (e.g. for sample exploration or when average lifetime information is sufficient), while information-rich TTTR data can be collected separately for more advanced post-processing methods including exponential fitting, phasor plot computation, and FCS-based algorithms.

In conclusion, in this work we present a simple and low-cost approach to implement accurate two-photon excited FLIM at state-of-the-art acquisition speeds in living animals, aiming at expanding the adoption of the technique among the scientific community and promoting a shift towards more quantitative microscopy studies.

## 5. Back matter

### 5.1 Funding

This work was funded by the Innosuisse Innovation Agency (Grant 52207.1).

## 5.2 Acknowledgement

We would like to thank Prof. Ivan Rech and Prof. Giulia Acconcia for the useful discussions.

## 5.4 Data availability statement

The code utilized to perform FLIM analysis is available at https://gitlab.com/einlabzurich/flimanalysis. The code to perform data acquisition is available upon request from the authors.

