## Supplementary Figures 1-6 for "A low-cost FPGA-based approach for pile-up corrected high-speed in vivo FLIM imaging"

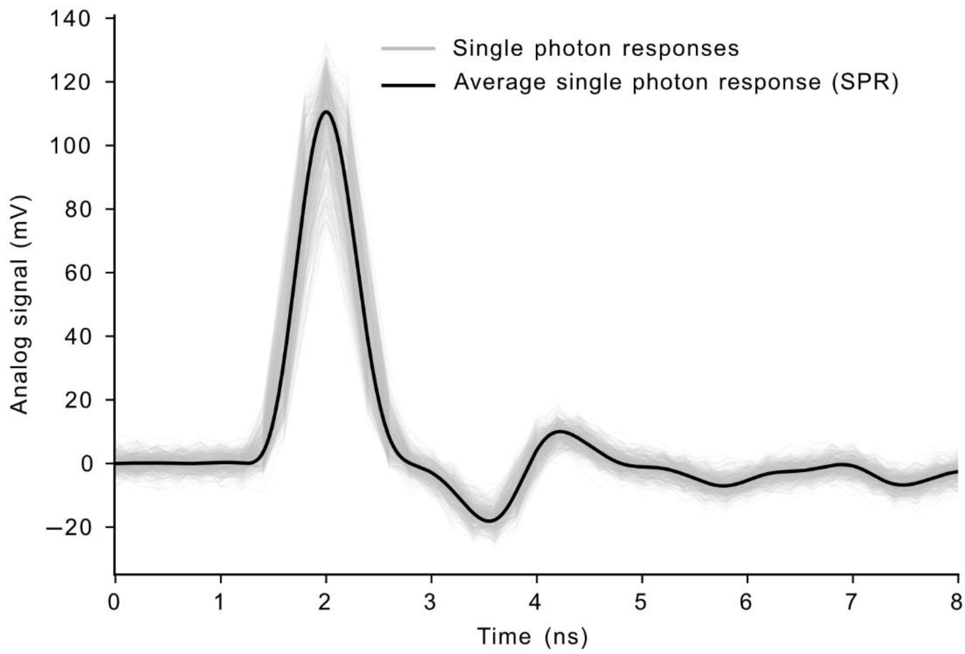


**Supplementary Figure 1:** Single-photon response of the PMA-40mod Hybrid Photodetector (PicoQuant, Germany) used in this study.

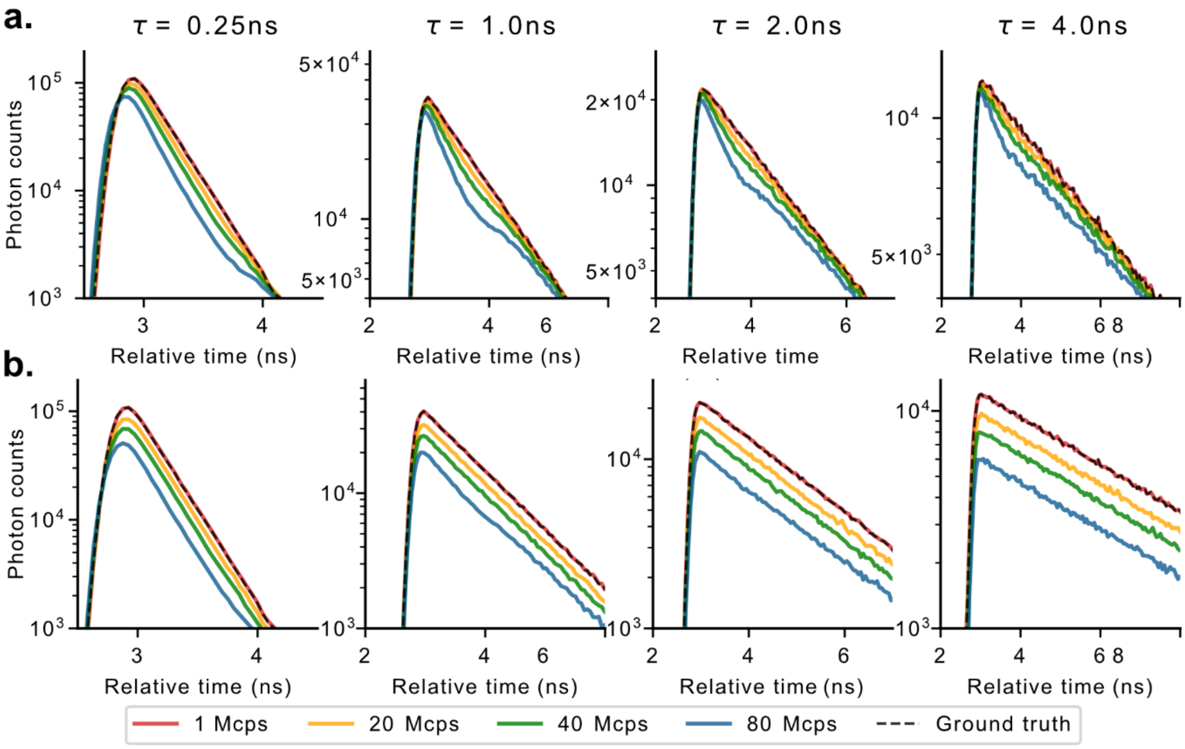


**Supplementary Figure 2:** Simulated decays for different monoexponential lifetimes and count rates with the before (a) and after (b) application of the LPBT correction. The SPR shown in Supplementary Figure 1 was used to simulate the detector response.

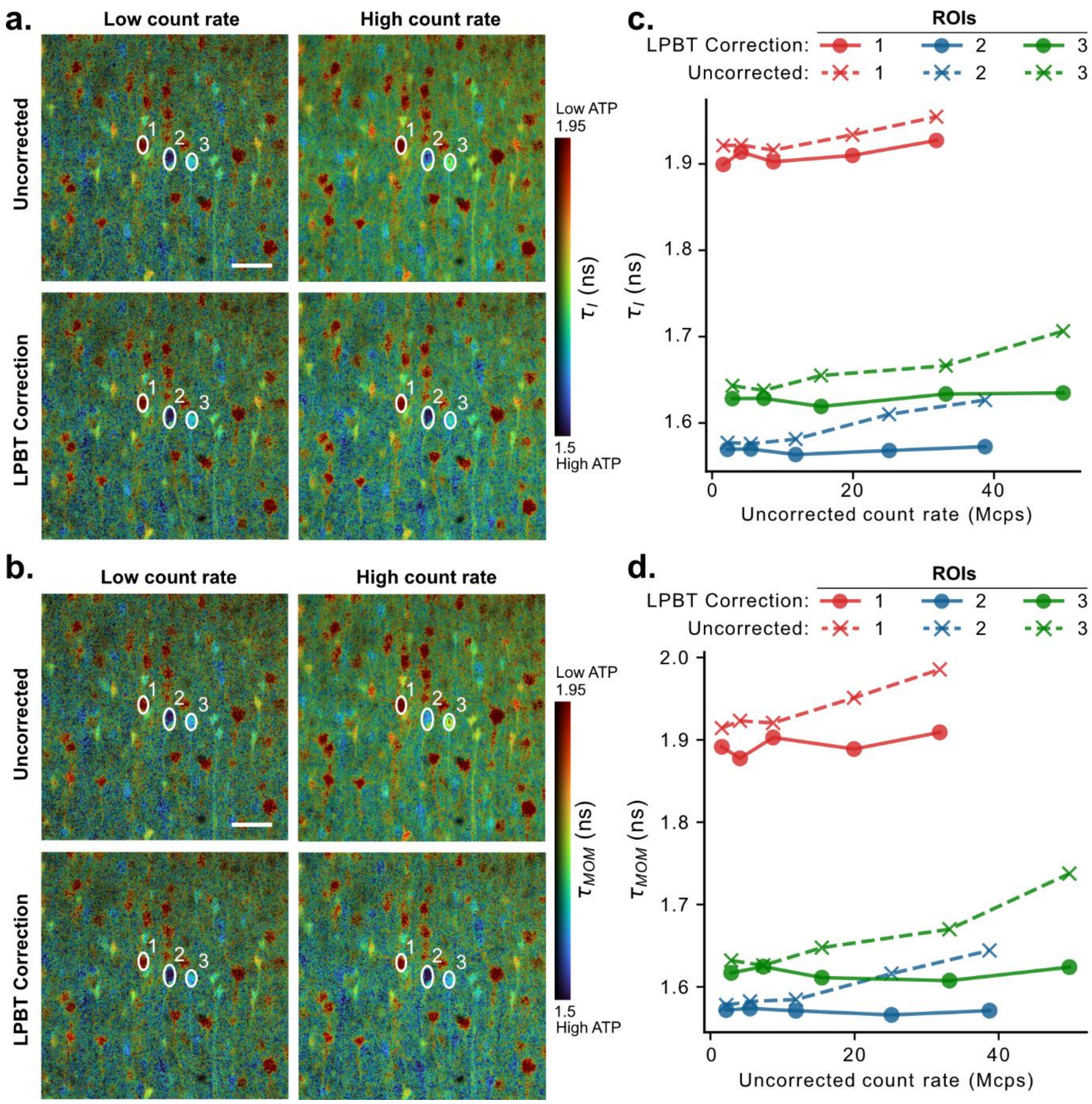


**Supplementary Figure 3:** (a,b) Comparison of LPBT corrected and uncorrected (detected) ATP lifetime images at low (<6 Mcps) and high (up to 85 Mcps) count rates and using different lifetime estimators: (a) $\tau_I$ and (b) $\tau_{MOM}$. (c, d) The change in $\tau_I$ (c) and $\tau_{MOM}$ (d) for each selected ROI as the detected count rate increases. Scale bar = 40 $\mu m$.

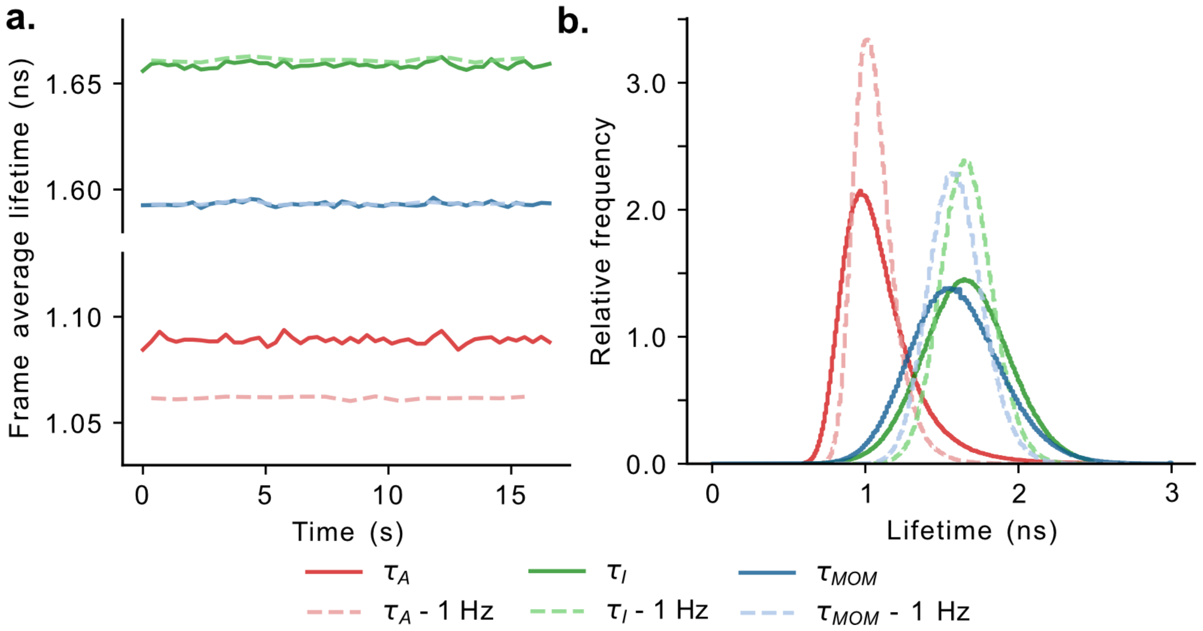


**Supplementary Figure 4:** (a) Frame average lifetime at baseline for ATP images using different estimators with and without time binning. (b) Distribution of pixel lifetime values at baseline using different estimators with and without time binning.

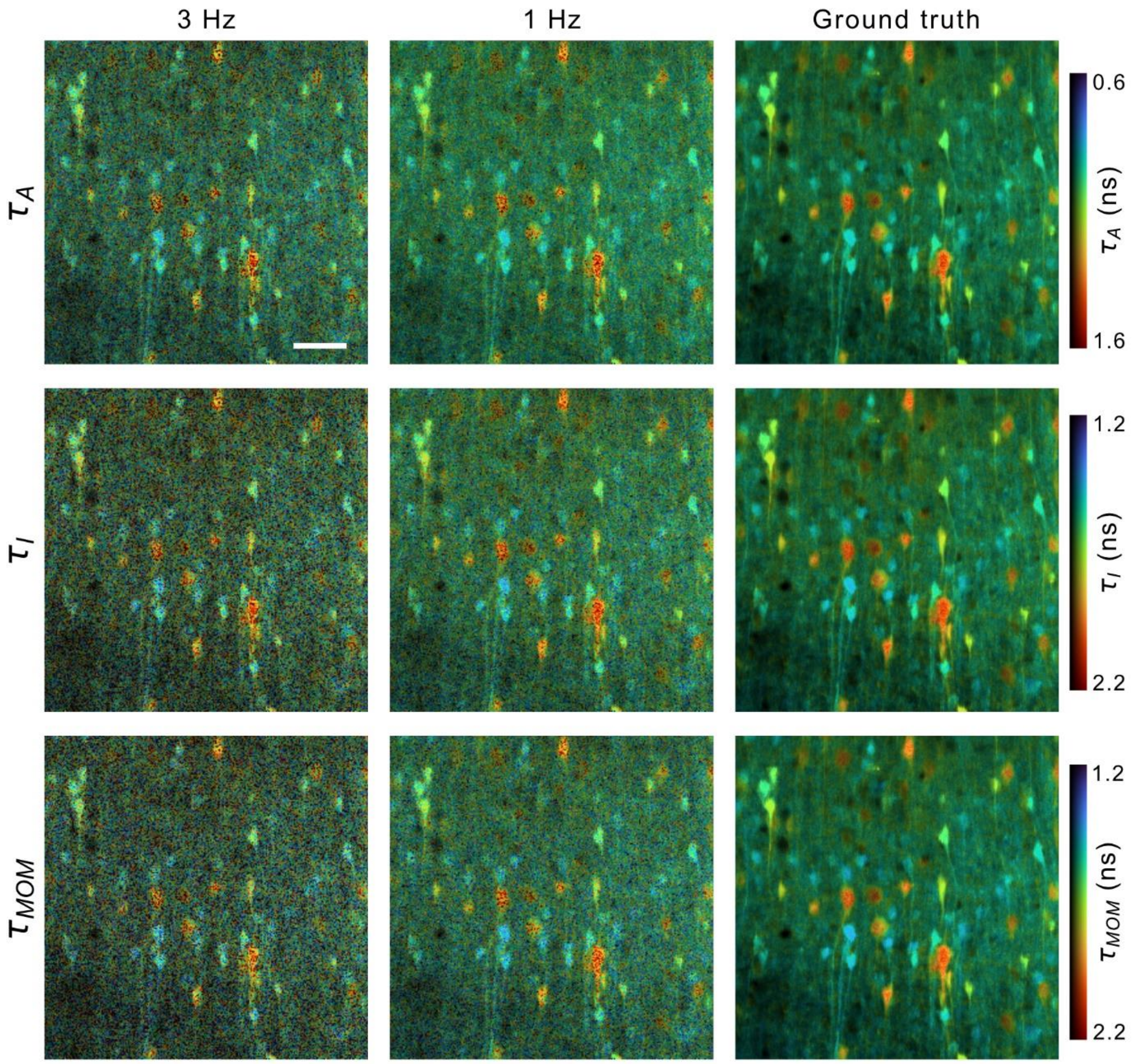


**Supplementary Figure 5:** Lifetime images of ATP at baseline for 3 Hz (single frame), 1 Hz (average of three frames), and ground truth (average of 150 frames) for all lifetime estimators at 256x256 pixels resolution. Scale bar = 40 $\mu m$.

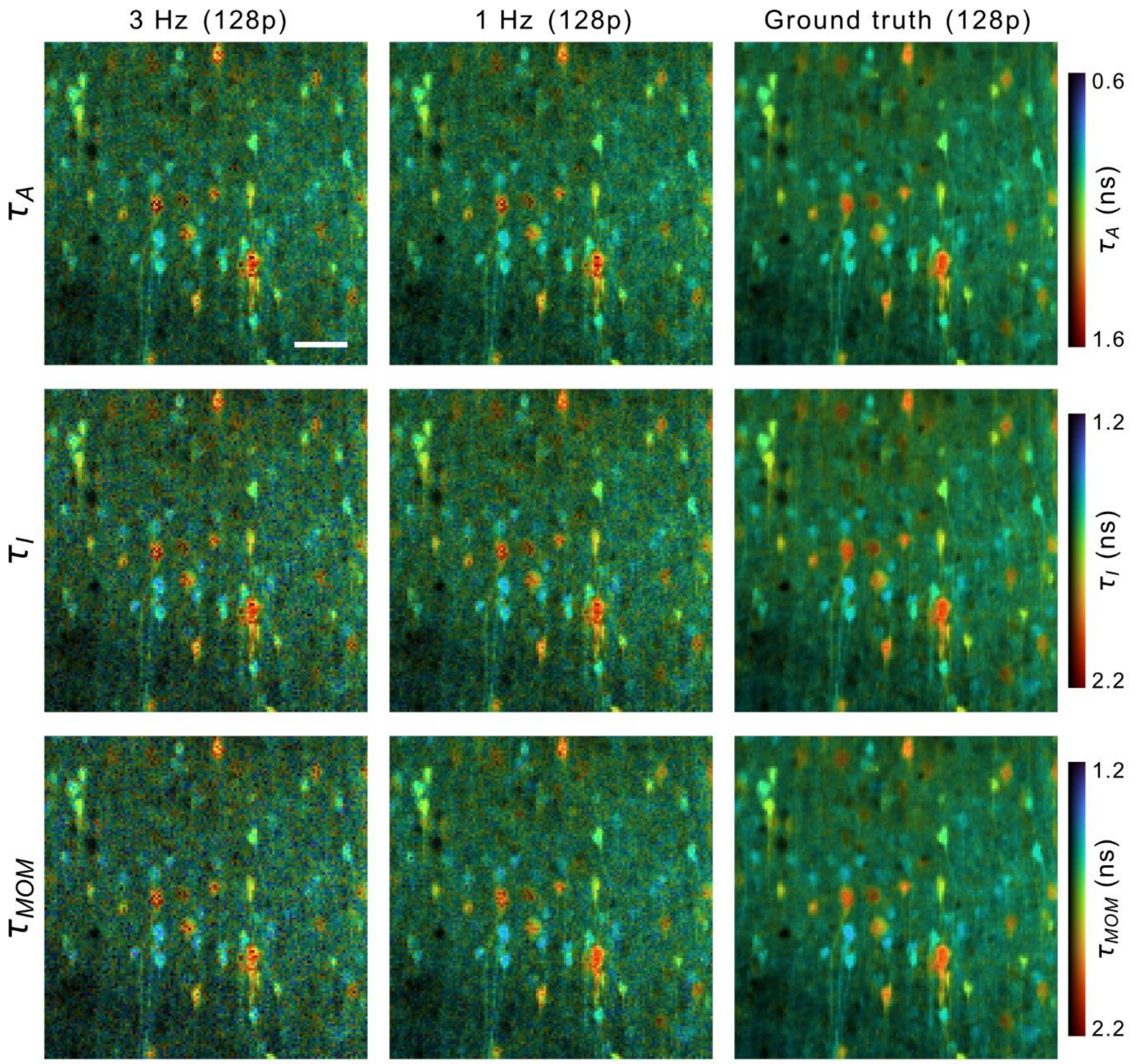


**Supplementary Figure 6:** Lifetime images of ATP at baseline for 3 Hz (single frame), 1 Hz (average of three frames), and ground truth (average of 150 frames) for all lifetime estimators at 128x128 pixels resolution. Scale bar = 40 $\mu m$.

**Supplementary Video 1:** Video recording showing the response of the ATP sensor Ateam1.03YEMK to $NaN_3$ application, at 256x256 resolution. The panel on the left shows the acquired FLIM data at the 3 Hz imaging speed while the panel on the right shows the result of applying temporal binning for every three frames, effectively reducing the imaging speed to 1 Hz. The plot on the bottom shows the method-of-moments lifetime ($\tau_{MOM}$) calculated as average across a frame at 3 Hz. To reduce the video length, the reproduction speed was increased 10 times with respect to the actual imaging speed. Scale bar = 40 $\mu m$.

**Supplementary Video 2:** Video recording showing the response of the ATP sensor Ateam1.03YEMK to $NaN_3$ application, at 128x128 resolution. The panel on the left shows the acquired FLIM data at the 3 Hz imaging speed while the panel on the right shows the result of applying temporal binning for every three frames, effectively reducing the imaging speed to 1 Hz. The plot on the bottom shows the method-of-moments lifetime ($\tau_{MOM}$) calculated as average across a frame at 3 Hz. To reduce the video length, the reproduction speed was increased 10 times with respect to the actual imaging speed. Scale bar = 40 $\mu m$.
